# Novel locus influencing retinal venular tortuosity is also associated with risk of coronary artery disease

**DOI:** 10.1101/121012

**Authors:** Abirami Veluchamy, Lucia Ballerini, Veronique Vitart, Katharina E Schraut, Mirna Kirin, Harry Campbell, Peter K Joshi, Devanjali Relan, Sarah Harris, Ellie Brown, Suraj K Vaidya, Bal Dhillon, Kaixin Zhou, Ewan R Pearson, Caroline Hayward, Ozren Polasek, Ian J Deary, Thomas MacGillivray, James F Wilson, Emanuele Trucco, Colin NA Palmer, Alexander S F Doney

## Abstract

Structural variation in retinal blood vessels is associated with global vascular health in humans and may provide a readily accessible indicator of several diseases of vascular origin. Increasing evidence suggests variation in retinal vasculature is highly heritable. This study aimed to identify genetic determinants of retinal vascular traits. We reported a meta-analysis of genome-wide association studies (GWAS) for quantitative retinal vascular traits derived using semi-automatic image analysis of digital retinal photographs from the Genetics of Diabetes Audit and Research in Tayside (GoDARTS) (n=1736) and the Orkney Complex Disease Study (ORCADES) (n=1358) cohorts. We identified a novel genome-wide significant locus at 19q13 (*ACTN4/CAPN12*) for retinal venular tortuosity (*TortV*), and one at 13q34 (*COL4A2*) for retinal arteriolar tortuosity (*TortA*); these two loci were subsequently confirmed in three independent cohorts (n=1413). In the combined analysis in *ACTN4/CAPN12* the lead single nucleotide polymorphism (SNP) was rs1808382 (n=4507; Beta=−0.109; standard error (SE) =0.015; P=2.39×10^−13^) and in *COL4A2* it was rs7991229 (n=4507; Beta=0.103; SE=0.015; P=4.66×10^−12^). Notably, the *ACTN4/CAPN12* locus associated with retinal *TortV* is also associated with coronary artery disease and heart rate. Our findings demonstrate the contribution of genetics in retinal tortuosity traits, and provide new insights into cardiovascular diseases.

**Author Summary:** Retinal vascular features are associated with wide range of diseases related to vascular health and provide an opportunity to understand early structural changes in vasculature which may help to predict disease risk. Emerging evidence indicates that retinal tortuosity traits are both associated with vascular health and highly heritable. However, the genetic architecture of retinal vascular tortuosity has not been investigated. We therefore performed a genome-wide association study on retinal arteriolar tortuosity (*TortA*) and retinal venular tortuosity trait (*TortV*) using data from two independent discovery cohorts of 3094 individuals of European-heritage. We found a novel associations at 19q13 (*ACTN4/CAPN12*) for *TortV*, and one at 13q34 (*COL4A2*) for *TortA* at discovery stage and validated in three independent cohorts. A significant association was subsequently found between lead SNPs at 19q13 and coronary artery disease, cardiovascular vascular risk factors and heart rate. We also performed genome-wide association studies for retinal vascular calibres and optic disc radius (*ODradius*) and replicated previously reported locus at 10q21.3 for *ODradius*. Our findings highlight genetic impacts on retinal venular tortuosity and it is association with cardiovascular disease. This may provide a molecular pathophysiological foundation for use of retinal vascular traits as biomarkers for cardiovascular diseases.

## Introduction

Retinal vascular traits can be readily measured non-invasively from fundus images and changes in these traits have been linked to a number of clinical conditions associated with vascular health including cardiovascular disease[1,2], stroke[3], hypertension[4], and neurodegenerative disease[5]. The association between retinal vascular calibers and cardiovascular disease has been reported in numerous studies and structural variation in retinal vasculature could predict cardiovascular risk[6–8]. More recently deep learning applied to retinal images has been successfully used to predict cardiovascular risk factors and outcomes[9].

Increasing evidence has shown a significant genetic component to variation in retinal blood vascular traits[10,11]. Understanding the molecular genetic architecture of retinal vascular features provides a molecular pathophysiological basis linking retinal microvascular features with systemic vascular pathology. Recent genome-wide association studies (GWAS) found number of loci for widely investigated retinal traits including central retinal vein equivalent (*CRVE*)[12–14], retinal arteriolar equivalent (*CRAE*)[12–14], and optic disc morphology[15]. Evidence suggests retinal vascular tortuosity, another potentially important vascular parameter, is also associated with a range of cardiovascular risk factors [16,17]. Heritability estimates for retinal arterial tortuosity range from 50-82% and 21% for retinal venular tortuosity[18,19], indicating a substantial genetic contribution to the variation in these parameters.

To our knowledge, no studies have performed genome-wide scan on retinal vascular tortuosity traits. We therefore carried out a discovery stage GWAS analysis using two independent cohorts including the Genetics of Diabetes Audit and Research in Tayside Study (GoDARTS) and the Orkney Complex Disease Study (ORCADES) to examine the underlying genetic factors influencing the retinal vascular tortuosity traits derived from digital retinal photographs; arteriolar tortuosity (*TortA*), maximum *TortA* (*TortAmax*), venular tortuosity (*TortV*), and maximum *TortV*(*TortVmax*). We also conducted a GWAS of other previously investigated retinal vascular traits including *CRAE, CRVE*, Arteriole-to-Venule ratio (*AVR*), as well as Optic Disc radius (*ODradius*). We confirmed our findings for retinal tortuosity traits in three independent replication cohorts (Lothian Birth Cohort 1936 (LBC1936), CROATIA-Korčula, and CROATIA-Split). Moreover, we examined the relationship between these sentinel SNPs and cardiovascular risk factors using data from the Coronary Artery Disease Genome wide Replication and meta-analysis plus The Coronary Artery Disease (CARDIoGRAMplus C4D) consortium meta-analysis[20], Global Lipid Genetics Consortium analysis[21] (GLGC), International consortium for blood pressure (ICBP) GWAS analysis[22] and on heart rate from the UK Biobank [23].

## Results

### Study samples

Two independent discovery cohorts were included; patients with type 2 diabetes from the GoDARTS (n=1736) and a population-based sample comprising the ORCADES (n=1358). In both these cohorts, traits were measured from retinal fundus images (**S1 Fig.**) using VAMPIRE 3.1 [24,25] (Vascular Assessment and Measurement Platform for Images of Retina), which enables efficient, semi-automatic measurement of the retinal vasculature from large numbers of images. The VAMPIRE methodology used in the discovery stage has been previously reported. [25–27]. The study design and characteristics of the discovery cohorts are shown in Fig 1., and Table 1.

**Table 1.** Descriptive statistics of variables for discovery study cohorts.

| Traits | GoDARTS | ORCADES |
| --- | --- | --- |
| Sample size | 1742 | 1358 |
| Age (yr.) | 69.78 ± 9.63 | 52.23 ± 14.7 |
| Male/Female | 990/752 | 542/816 |
| *lnTortA* (mean ± SD) | -9.76 ± 0.99 | -10.94 ± 1.32 |
| *lnTortA* max (mean ± SD) | -8.41 ± 0.92 | -9.22 ± 1.17 |
| *lnTortV* (mean ± SD) | -10.03 ± 0.78 | -11.52 ± 1.01 |
| *lnTortV* max (mean ± SD) | -8.66 ± 0.86 | -9.76 ± 0.90 |
| ODradius (mean ± SD) | 208.56 ± 18.97 | 183.26 ± 19.46 |
| CRAE (mean ± SD) | 33.11 ± 2.39 | 29.09 ± 2.55 |
| CRVE (mean ± SD) | 43.74 ± 3.11 | 39.15 ± 3.62 |
| AVR (mean ± SD) | 0.76 ± 0.05 | 0.75 ± 0.08 |

Eight retinal vascular parameters are abbreviated as follows: Natural log transformed – Retinal Arteriolar Tortuosity (*lnTortA*), maximum TortA (*lnTortAmax*), Retinal Venular Tortuosity (*lnTortV*), and maximum TortV (*lnTortV*). Optic Disc radius (ODradius), Central Retinal Arteriolar Equivalent (CRAE), Arteriole-to-Venule ratio (AVR), Central Retinal Venular Equivalent (CRVE), SD – standard deviation. GoDARTS: Genetics of Diabetes Audit and Research in Tayside; ORCADES: Orkney Complex Disease Study.

**Fig 1.**
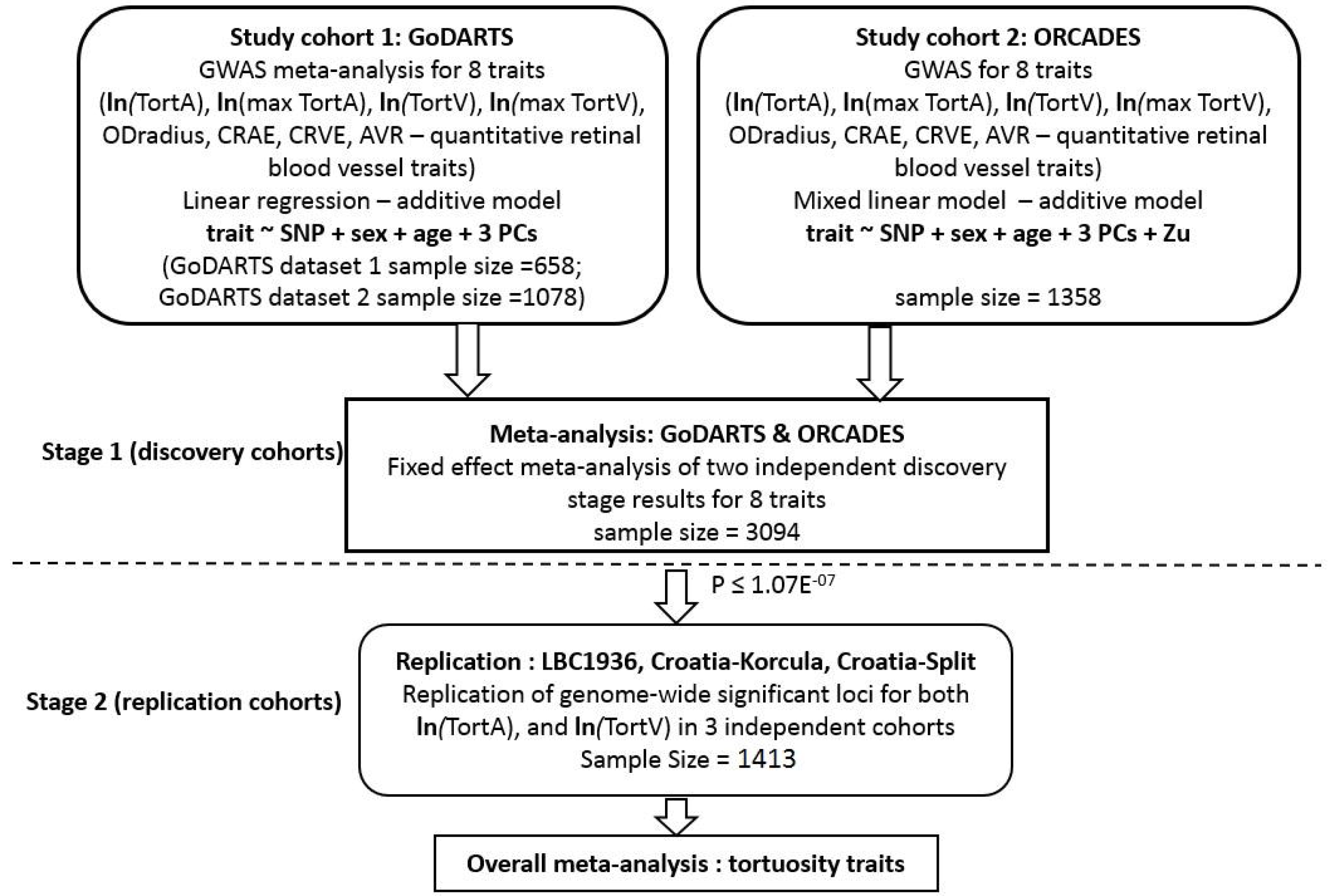
Study Design. GoDARTS: Genetics of Diabetes Audit and Research in Tayside; ORCADES: Orkney Complex Disease Study; LBC1936: Lothian Birth Cohorts 1936; All Croatia: Croatia island of Korcula, Croatia Split; *ODradius*: Optic Disc Radius, *CRAE*: Central Retinal Arteriolar Equivalent, *CRVE*: Central Retinal Venular Equivalent, *AVR*: Arteriole-to-Venule ratio, Natural log transformed data – *TortA*: retinal arteriolar tortuosity, *TortAmax*. maximum retinal arteriolar tortuosity, *TortV*: retinal venular tortuosity, *TortVmax*. maximum retinal arteriolar tortuosity; PC: Principal Components; u is the genetic value for each subject under a random effects model, covariance amongst subjects assumed to be proportionate to the genomic relationship matrix.

In the discovery stage, we performed a GWAS using the GoDARTS cohort for each retinal trait separately and tested the additive effect of each variant, adjusted for age, gender and the first three principal components. Similarly, GWAS was performed for the same traits in the ORCADES cohort, using a mixed model to account for kinship and first three principal components as covariates correcting for population structure. As there were statistically significant differences by age, and gender between two discovery cohorts (both age and gender, P<0.0001) the model was adjusted for these in both discovery cohorts. We combined the summary results from these two cohorts for each trait using a fixed effect meta-analysis and the genomic inflation factor is 0.99. **S1 Table** presents the results from the meta-analysis and independent cohort GWAS analysis. Manhattan plots and regions of interest for tortuosity traits are shown in Fig 2–3. Manhattan plots for other retinal traits and QQ plots for all traits are shown in, and **S2-S4 Figs**.

**Fig 2.**
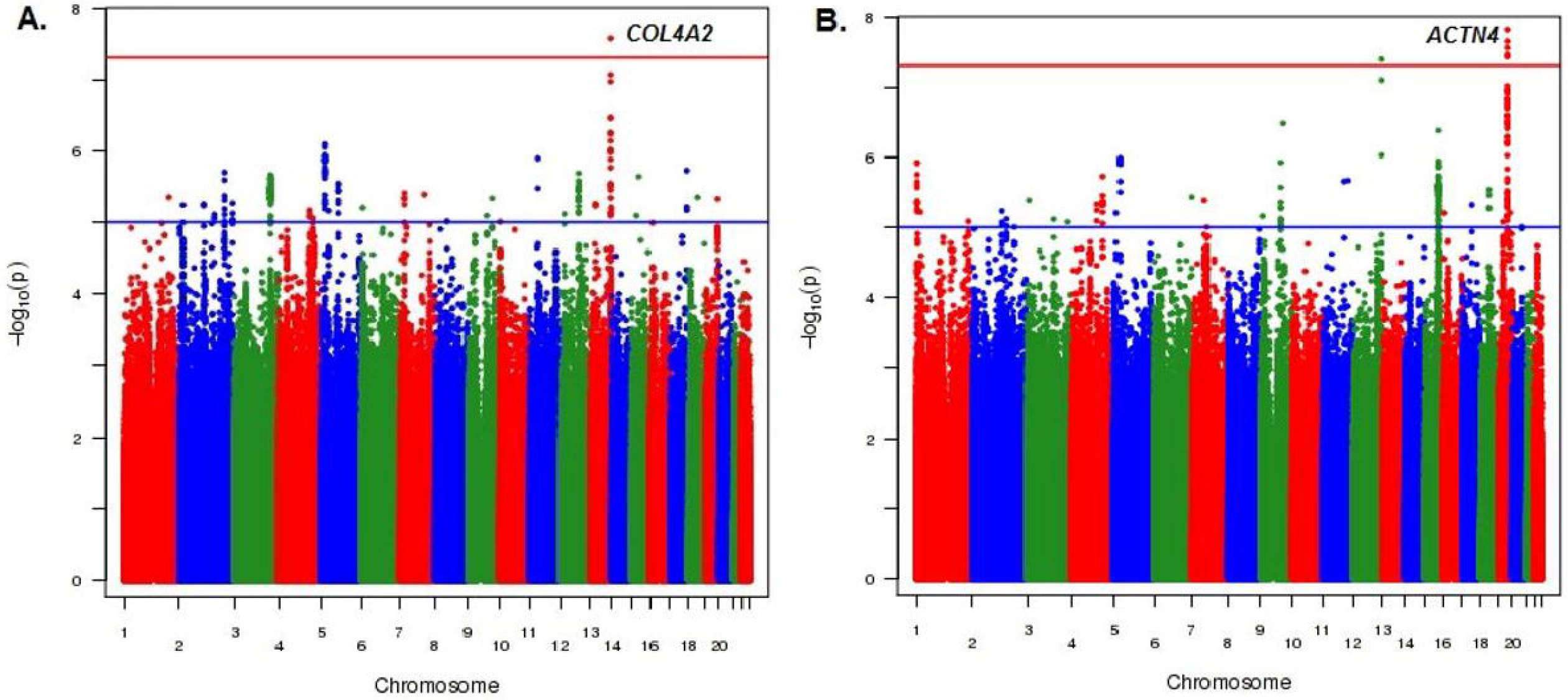
Manhattan plots for meta-analysis of genome-wide association results from two independent discovery cohorts. A, represents the results for the arteriolar tortuosity (*TortA*) and B. represents the results for the venular tortuosity trait (*TortV*). The blue and red horizontal lines indicate the suggestive and genome-wide significance threshold (P<5×10^−8^), respectively.

**Fig 3.**
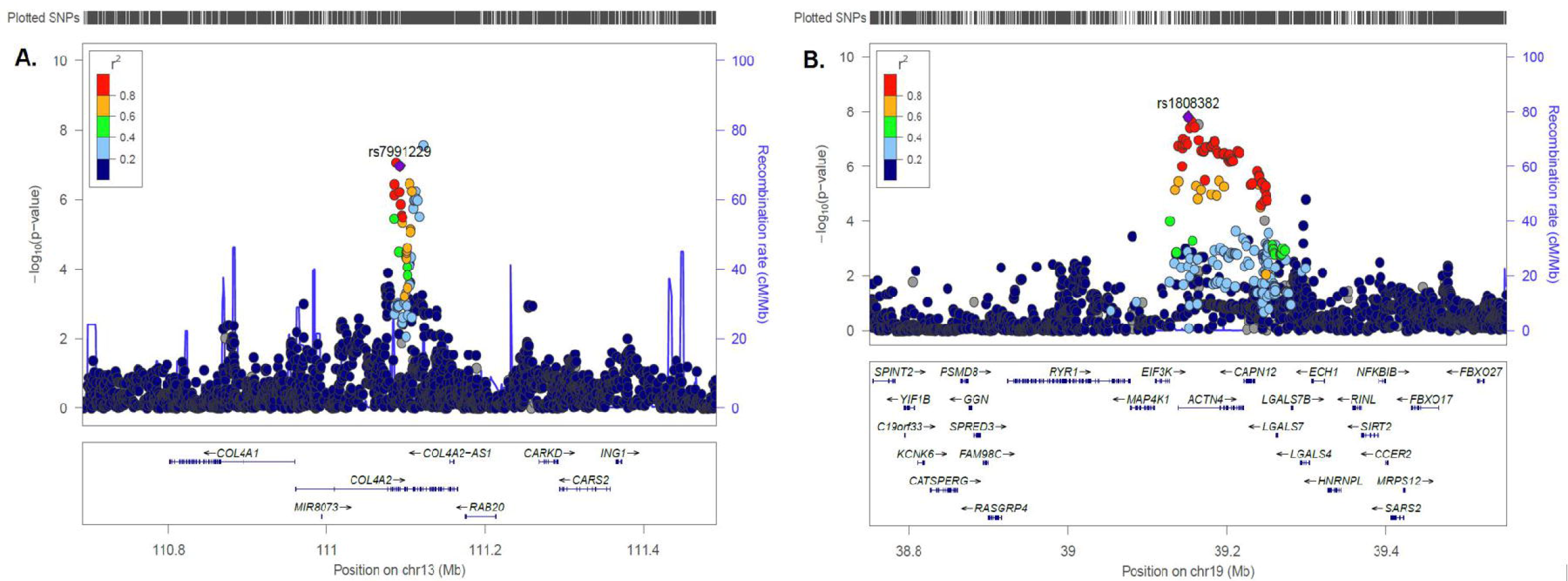
Regional association and recombination plots of variants that reached genome-wide significance in overall meta-analysis (discovery and replication stage). A-B, Top hits for *TortA* and C-D, Top hits for *TortV*. Each plot was created using LocusZoom for the lead SNP in genomic region 400 kb in either side of the significant signal. Blue spikes represents the estimated recombination rates. Colour scale (high to low r2) circles depicts the pairwise correlation (r2) between lead SNP and other SNPs in the loci. The lead SNP in that region is indicated by purple colour solid diamond and gene annotations in this region is shown in the bottom panels.

This analysis revealed one genome-wide significant (p<5×10^−8^) SNP, rs56399312, associated with *TortA* at 13q34, in *COL4A2* with moderate heterogeneity (I^2^=0.50); Beta=0.182, SE= 0.032, P= 2.70 ×10^−8^, and another SNP rs9515212 near *COL4A2* that was just below the threshold for genome-wide significance; Beta=0.151, SE= 0.028, P= 8.59 ×10^−8^. Conditional analysis on the lead SNP indicated that these are not independent signals (**S2 Table**). Two genome-wide significant SNPs were associated with *TortV*, at 19q13 in *ACTN4* (lead SNP rs1808382; Beta=−0.123, SE= 0.022, P= 1.55 ×10^−8^; no heterogeneity, I^2^=0.00), and at 12q24.33 near *TMEM132D* (lead SNP rs73157566; Beta= −0.294, SE= 0.054, P= 4.07 ×10^−8^; low heterogeneity, I^2^=0.10); these associations at both these loci have not been reported previously with any retinal vascular parameters.

Although we replicated previously reported loci for *CRVE*, we did not find any novel genome-wide significant loci for this trait[12]’[14]. In addition, we did not replicate any of the previously reported SNPs associated with *CRAE*[13]. Finally, we replicated a previously reported genome-wide significant locus for *ODradius* at 10q21.3 near *PBLD/ATOH7* (lead SNP rs61854835; Beta= −3.840, SE= 0.575, P= 4.06 ×10^−11^) and confirmed a number of other loci for this trait [15,28–30] (**S3 Table**).

We selected three lead SNPs near *ACTN4, TMEM132D*, and *COL4A2*, that reached significance P ≤ 1.07×10^−07^ as well as their effect size and direction being similar across the discovery cohorts, as candidates to carry forward for replication, and confirmed these in three independent cohorts comprised of up to 1413 individuals of European ancestry including the Lothian Birth Cohort 1936[31] (LBC1936), Croatia-Korcula, and Croatia-Split. Retinal images from these cohorts had been analyzed by SIVA 3.1 [32,33] (Singapore I Vessels Assessment) software to quantify the tortuosity traits. The characteristics of the replication cohorts have been presented in Table 2. Two *TortA*-associated SNPs, rs7991229, and rs9515212 in *COL4A2* reached significance (P<0.05) in the LBC1936 and Croatians cohorts but lead SNP (rs56399312) did not replicate in the LBC1936. Two TortV-associated SNPs, rs1808382 and rs3786835 in *ACTN4/CAPN12* reached suggestive significance (P<1×10^−04^) in the combined analysis of replication cohorts whereas rs73157566 near *TMEM132D* did not replicate.

**Table 2.**
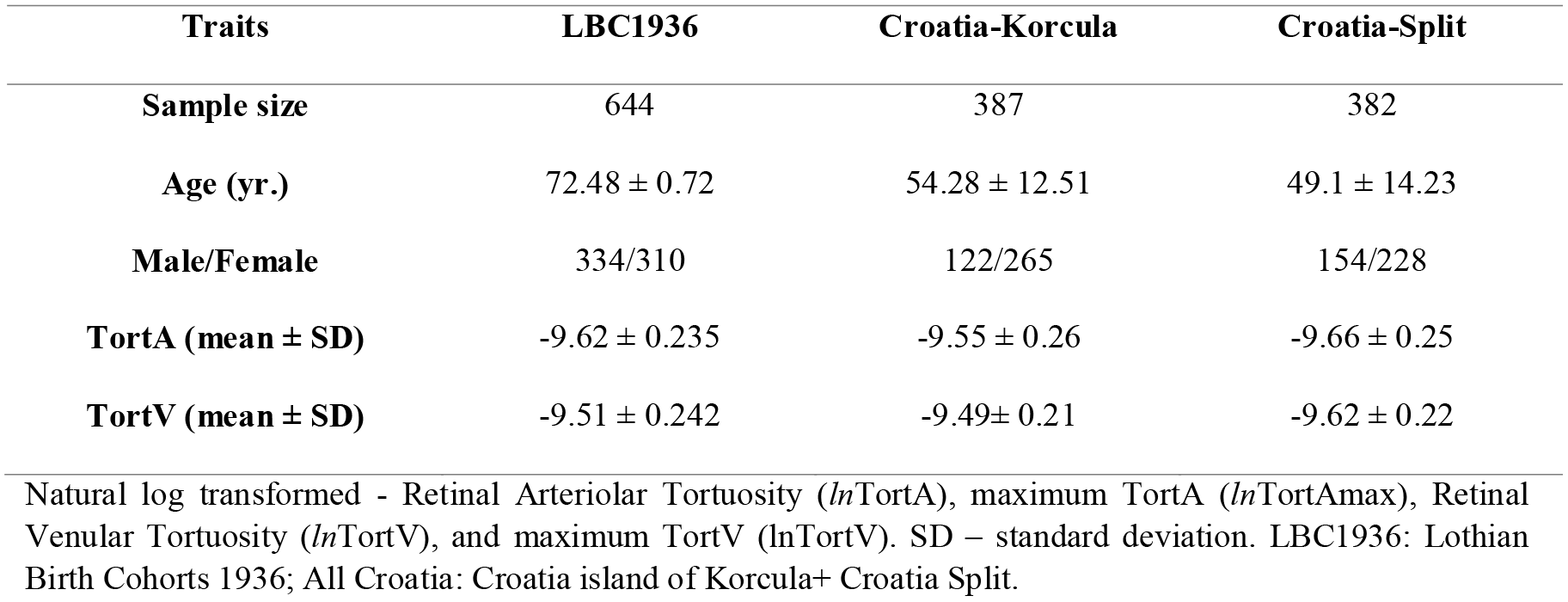
Descriptive statistics of variables for replication study cohorts.

| Traits | LBC1936 | Croatia-Korcula | Croatia-Split |
| --- | --- | --- | --- |
| Sample size | 644 | 387 | 382 |
| Age (yr.) | 72.48 ± 0.72 | 54.28 ± 12.51 | 49.1 ± 14.23 |
| Male/Female | 334/310 | 122/265 | 154/228 |
| TortA (mean ± SD) | -9.62 ± 0.235 | -9.55 ± 0.26 | -9.66 ± 0.25 |
| TortV (mean ± SD) | -9.51 ± 0.242 | -9.49 ± 0.21 | -9.62 ± 0.22 |

Natural log transformed - Retinal Arteriolar Tortuosity (*lnTortA*), maximum TortA (*lnTortAmax*), Retinal Venular Tortuosity (*lnTortV*), and maximum TortV (*lnTortVmax*). SD – standard deviation. LBC1936: Lothian Birth Cohorts 1936; All Croatia: Croatia island of Korcula+ Croatia Split.

### Meta-analysis of discovery and replication cohorts

In the overall meta-analysis, only SNPs at 13q34, *COL4A2* (*TortA*) and 19q13, *ACTN4* (*TortV*) were confirmed at genome-wide significance. Although *TortA* associated SNPs rs9515212 and rs7991229 were not genome-wide significant in the discovery meta-analysis, they reached genome-wide significance in the overall meta-analysis and no heterogeneity (I^2^=0.00) was observed across different cohorts; Poverall=4.66×10^−12^ and P_overall_=4.71×10^−12^, respectively. Whereas the lead SNP in *COL4A2* for *TortA* (rs56399312) at discovery stage did not reach genome-wide significance in overall meta-analysis (P_overall_=1.95×10^−07^). For *TortV* the lead SNPs, rs1808382 (P_overall_=2.39×10^−13^) and rs3786835 (P_overall_=3.31×10^−13^) near *ACTN4/CAPN12*, maintained genome-wide significance with no heterogeneity (I^2^ =0.00). These SNPs are in tight LD and therefore do not represent independent signals. Table 3 contains the summary statistics from replication cohorts and meta-analysis of these cohorts. Forest plots for lead SNPs in the combined analysis are shown in Fig 4.

**Fig 4.**
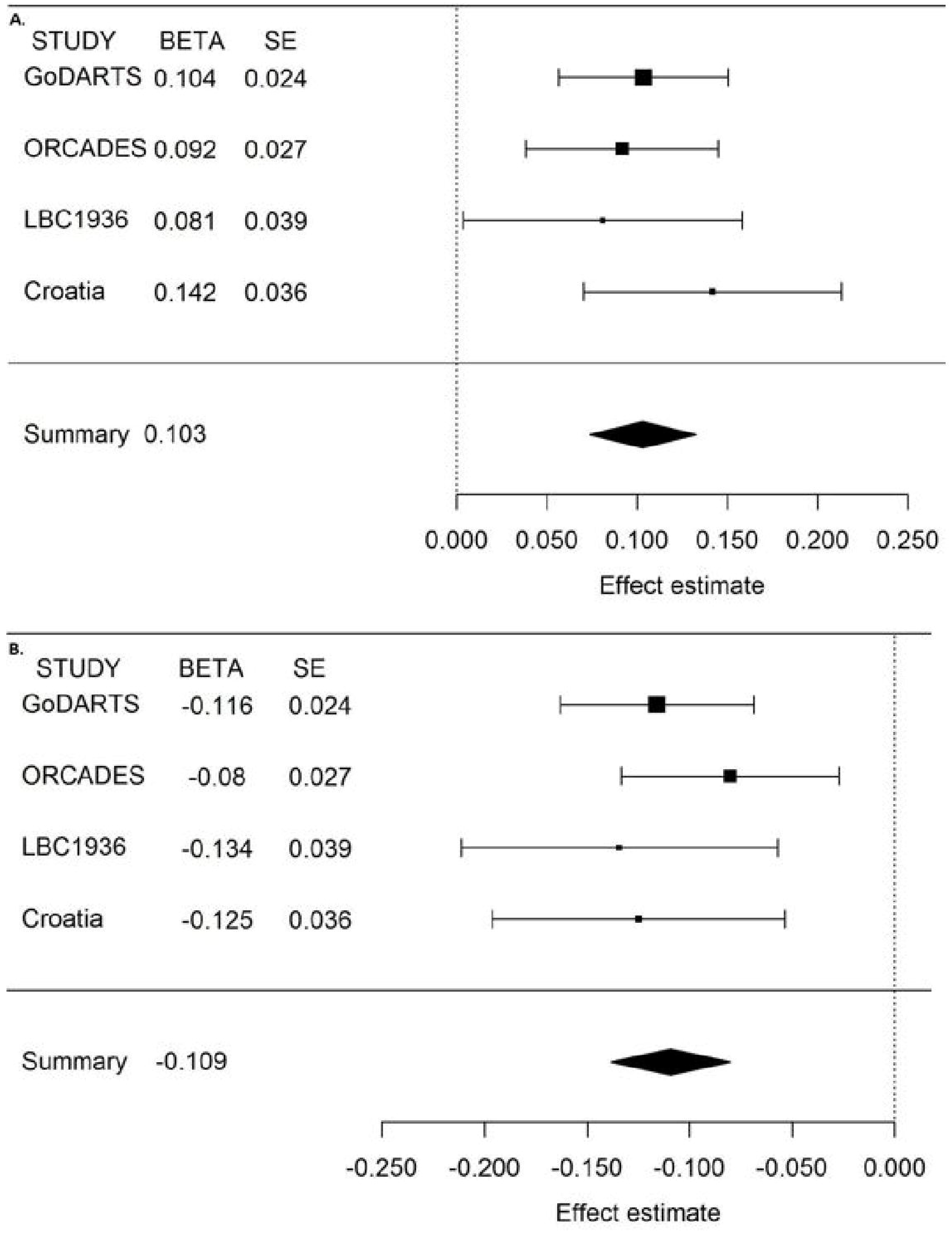
Forest Plots for the genome-wide significant hits (overall meta-analysis) associated with arteriolar (A. rs7991229) and venular tortuosity traits (B. rs1808382). The plots represent standardized beta and standard error from GoDARTS, ORCADES, LBC1936, Croatia KORCULA-SPLIT, and meta-analysis study. Standardized beta estimate: Change in natural log transformed retinal tortuosity traits for each copy of the effect allele. Due to the difference in the units of the beta and standard errors between the discovery and replication studies arising from different approaches to measurement, we standardized the effect estimates from each individual’s study results.

### *TortA*-associated variants

*COL4A2* encodes collagen type IV alpha 2, one of the six subunits of type IV collagens which are major structural components of basement membranes, forming a thin sheet of fibers under the endothelium controlling passage of vasoactive substances. These are conserved across species and C-terminal non-collagenous domains play a role in angiogenesis[34]. Recent GWAS report that common variants around *COL4A2* and *COL4A1* (a paralogue immediately proximal to *COL4A2*, with which it shares a promoter and is coexpressed), are associated with coronary artery calcification[35], arterial stiffness [36], and coronary artery disease[20,37–39] (CAD).

**Table 3.**
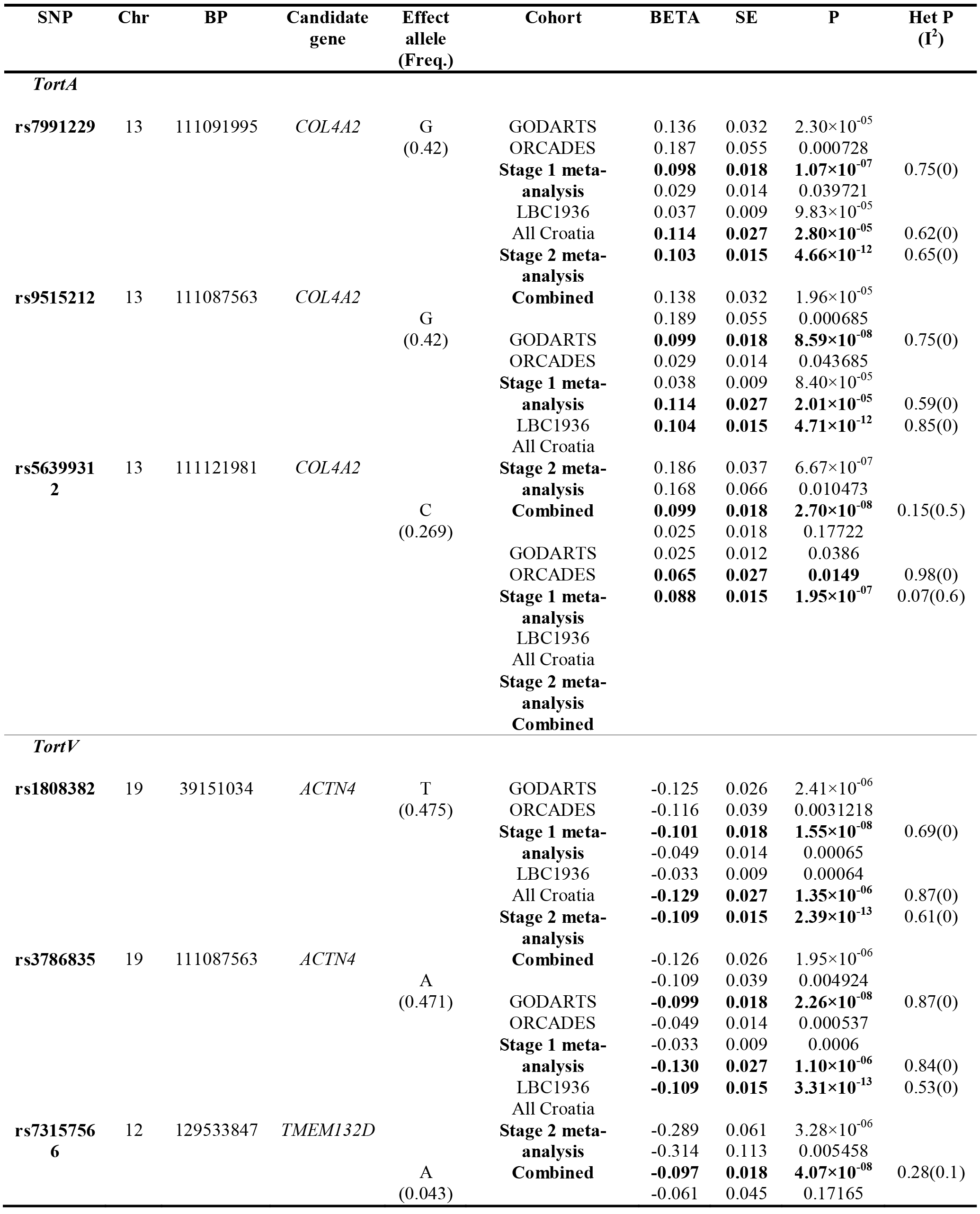
Results of discovery, replication and overall meta-analysis for tortuosity traits.

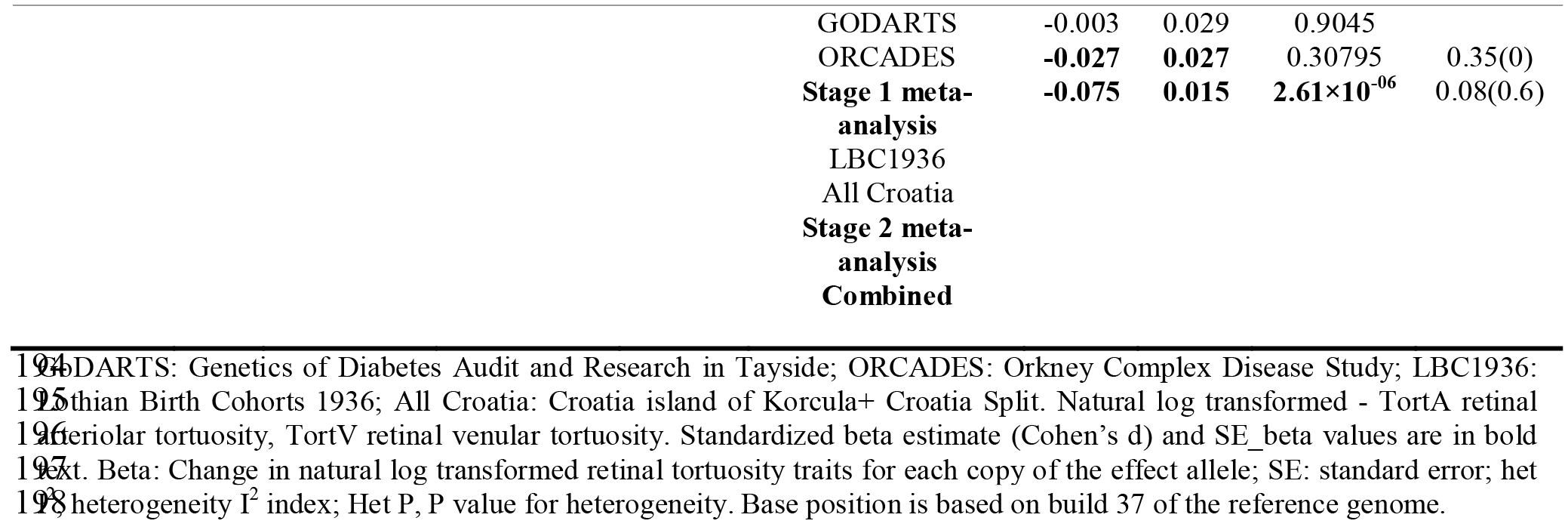

Interestingly, gene expression data from Gene Atlas [40], a human protein-coding transcriptome study validated the high expression of *COL4A2* in retinal micro-vessel endothelial cells (**S5 Fig.**) whereas *COL4A1* is weakly expressed in retina indicating a specific role of *COL4A2* in the retinal vasculature. TortA-associated variants near *COL4A2* significantly alter transcription factor binding motifs and have putative effects on transcription as annotated by ENCODE (**S4 Table**). Additionally, expression data from the GTEx database[41] confirmed that these significant SNPs, are associated with the expression of *COL4A2* in heart left ventricle and artery aorta, shown in **S5 Table**, **S6 Fig.**, and these SNPs are in linkage disequilibrium (LD; r^2^=0.99, D’=1).

Lead SNPs associated with *TortA* remained significant after conditioning on the previously reported cardiovascular risk variants in *COL4A2* (rs11617955[20], rs4773144[38], rs9515203[39]) (**S7 Fig.**, **S6 Table**). Conversely the lead SNPs for *TortA* were not associated with coronary artery disease (CAD) and myocardial infarction (MI) risk in the CARDIoGRAMplus C4D consortium meta-analysis[20] (**S7 Table**). Finally, the CAD associated variants specifically in *COL4A1* from the CARDIOGRAMplusC4D were not associated with *TortA*, whereas CAD associated *COL4A2* variants are only weakly associated with *TortA* (**S7-S8 Table**). Retinal vascular tortuosity traits have been previously associated with blood pressure[17,19] which may be therefore link these variants with CAD, however, we found no evidence for an association between these lead variants and blood pressure in the ICBP GWAS analysis[22] (**S9 Table**).

### *TortV*-associated variants

*ACTN4* encodes alpha-actinin 4, a cross-linking protein belonging to the spectrin superfamily and mutations in this gene cause focal segmental glomerulosclerosis in humans. *ACTN2*, a homolog of *ACTN4*, interacts with *ACTN4* and missense mutations in *ACTN2* are linked to a range of cardiac diseases[42]. Annotation by ENCODE[43] indicates that the two genome-wide significant variants (rs1808382, rs3786835) associated with *TortV* near *ACTN4* may have direct regulatory effects as they are located within a DNase I hypersensitivity site and in genomic regions enriched for promoter/enhancer histone marks in heart tissues (**S5 Table**). *ACTN4* and *CAPN12* (calcium-activated neural proteases) overlap by 339 bases at their 3’ ends and multi-tissue expression quantitative trait loci (eQTL) analysis confirms that these SNPs in *ACTN4* are associated with mRNA expression of both *ACTN4* and *CAPN12* in aorta, tibial artery, atrial appendage and left ventricle of the heart (**S5 Table**, **S8 Fig.**). Additionally, this analysis indicates that the T allele at rs1808382 is correlated with lower *ACTN4* (artery aorta; P=2.1×10^−03^) and this correlation is even stronger with *CAPN12* (artery aorta; P=2.0×10^−07^). However, while gene expression data using GENEINVESTIGATOR validated the high expression of *ACTN4* in arterial tissue, the highest expression of *CAPN12* appears to be in the hematopoietic system (**S5 Fig.**).

Lead SNPs in *ACTN4* were significantly associated with coronary artery disease in the CARDIoGRAMplus C4D consortium meta-analysis[20] (**S7 Table**) and were associated with CAD risk factors; HDL cholesterol and triglycerides in the GLGC[21], but not associated with blood pressure in the ICBP GWAS analysis[22] (**S9 Table**). Furthermore, we have confirmed the association between *TortV* and top variants in *ACTN4/CAPN12* in a sensitivity analysis that only included GoDARTS samples without any cardiovascular events prior to the retinal screening date (**S10 Table**). Moreover, recent meta-analysis of 35 GWAS studies reported the association of SNP (rs11083475) in the *ACTN4* locus with increased resting heart rate[44] which may increase cardiovascular disease risk. This signal is the same as that for *TortV* with strong LD being observed between the lead SNPs for *TortV* and the index SNP for heart rate. Furthermore we found that these SNPs were associated with heart rate in UK Biobank[23] (**S11 Table**, **S9 Fig.**).

## Discussion

In this first GWAS for quantitative retinal vascular tortuosity traits, we found novel loci for retinal arteriolar tortuosity (*COL4A2*) and for retinal venular tortuosity (*ACTN4/CAPN12*), which were replicated in three independent cohorts. Our findings are consistent, non-heterogeneous and have the same direction of effects across all five fairly homogeneous cohorts of European ancestry comprising individuals with and without diabetes, and irrespective of the measurement platform used. Notably, we also identified a genome-wide significant signal at a previously reported locus in/near *ATOH7/PBLD* for the optic disc radius and replicated previously identified variants for *CRVE* which validate our retinal traits measurement methods. Together these aspects strongly support the robustness our study design and findings. Power calculations indicated that a sample size of 4507 (stage 1 and stage 2), and 3094 (stage 1) and the effect size of 0.10 using Bonferroni correction (P<5×10^−8^) was adequate to provide 80% statistical power to detect the associations.

Previous studies have reported the association between *COL4A2* and CAD but the TortA-associated variants in *COL4A2* in the present study are not associated with cardiovascular disease and similarly *COL4A2* variants that are associated with CAD do not appear to be associated with arteriolar tortuosity suggesting that variants in this gene complex may be involved differentially in the pathophysiology of microvascular and macrovascular diseases. However, more work has to be done to determine the distinct role of genetic variants in *COL4A2/COL4A1* in different clinical conditions. In contrast, we found that retinal venular tortuosity-associated variants were associated with coronary artery disease as well as heart rate. Furthermore, the lead variant influences the expression of the *ACTN4/CAPN12* genes in the heart tissue. Our sensitivity analyses including samples without CAD prior to the date of acquisition of the measured retinal image indicates that relationship between genetic predictors of retinal venular tortuosity and cardiovascular diseases is not due to reverse causation and demonstrate the robustness of our findings.

Evidence suggests that both retinal arteriolar and venular tortuosity traits are associated with blood pressure and cardiovascular risk factors[17]. However, a recent study from the ORCADES and Croatia-Korcula cohorts reported very weak association between retinal arteriolar tortuosity and blood pressure whereas no evident association between retinal venular tortuosity traits and blood pressure[19]. In this regard, neither the *TortA* nor *TortV-* associated variants were associated with systolic and diastolic blood pressure in the ICBP GWAS analysis thus it seems unlikely that observed associations with CAD or related traits are mediated through blood pressure.

In summary, this first GWAS for retinal arteriolar and venular tortuosity reveals SNPs influencing expression of *COL4A2* and *ACTN4/CAPN12* respectively. Our results demonstrate that the TortA-associated variants in *COL4A2* are independent of CAD, MI, and blood pressure, and point to a selective role of *COL4A2* rather than *COL4A1* in the retinal vessels. Strikingly, we found TortV-associated *ACTN4/CAPN12* SNPs are associated with CAD and heart rate but not associated with blood pressure. However, detailed investigation and functional validation of this new finding is essential to elucidate the causal roles of *ACTN4* and/or *CAPN12* in the observed cardiovascular pathophysiology. These findings highlight the potential genetic impacts of retinal vasculature to provide new insights into cardiovascular disease.

## Materials and Methods

### Study participants

#### Discovery Cohorts

Participants in the discovery phase of this study were obtained from the two independent cohorts, the GoDARTS [45] and the ORCADES. GoDARTS comprises individuals of European-heritage from Tayside, Scotland who provided a sample of blood for genetic analysis and consent to link their genetic information to the anonymized electronic health records. Approval for recruitment to GoDARTS was obtained from the Tayside Committee on Medical Research Ethics. 18,190 individuals were recruited with approximately half having type 2 diabetes at the time of recruitment with the other half being diabetes free. 7,290 individuals currently have genome-wide data for analysis. ORCADES is a family-based study of 2078 individuals aged 16-100 years recruited between 2005 and 2011 in the isolated Scottish archipelago of Orkney[46]. Genetic diversity in this population is decreased compared to Mainland Scotland, consistent with the high levels of endogamy historically. Fasting blood samples were collected and over 300 health-related phenotypes and environmental exposures were measured in each individual. All participants provided written informed consent and the study was approved by Research Ethics Committees in Orkney and Aberdeen.

#### Replication Cohorts

The LBC1936 comprises 1091 participants who were born in 1936, most of whom took part in the Scottish Mental Survey of 1947. At a mean age of 69.5 years (SD 0.8), between 2004 and 2007, they were recruited to a study to determine influences on cognitive aging[31]. The CROATIA-Korčula study comprises individuals from the Adriatic island of Korčula, between the ages of 18 and 88. The fieldwork was performed in 2007 in the eastern part of the island, targeting healthy volunteers who underwent complete eye examination and provided their blood sample for genetic analysis from the town of Korčula and the villages of Lumbarda, Žrnovo and Račišće. The Croatia-Split study included inhabitants of the Croatian coastal city of Split, aged 18 to 93. The sampling scheme was similar to Croatia-Korčula, and it took place during 2008 and 2009.

### Retinal Vascular Parameters Measurement

#### Retinal image analysis

##### Discovery cohorts

Standard digital retinal photographs used for routine diabetic retinopathy screening were obtained from the clinical record in 2,104 participants in GoDARTS. Images of the right eye of usable quality, defined using criteria reported in[25,26], were selected and categorized into two datasets based on the image pixel resolution: GoDARTS dataset 1 (n=788) and GoDARTS dataset 2 (n=1288). Finally, 661 images from the GoDARTS dataset 1, and 1083 images from GoDARTS dataset 2 were included after quality control (QC). 28 individual’s images from the GoDARTS were excluded due to inadequate resolution. Standard fundus retinal photographs centred between the macula and optic disc were obtained using digital fundus camera from 1,743 participants in ORCADES. After image processing and QC, 1595 individual’s retinal images were used for this study.

VAMPIRE 3.0, was used to measure retinal vascular traits in fundus images from both GoDARTS and ORCADES. The measurement process is organized as a sequence of automatic and manual stages. Manual stages allowed correction of errors made by the automatic software (e.g. vessel labeling as artery or vein) and to minimize their impact on statistical analysis. Standard protocols were followed to measure the retinal vessel parameters. Briefly, after automatic detection of the optic disc and its radius (*ODradius*), the 6 thickest arterioles and 6 thickest venules appearing in a zone extending out from the optic disc boundary to 2 optic disc diameters were sampled to calculate the median (*TortA*) and maximum (*TortAmax*) arteriolar tortuosity and the median (*TortV*) and maximum (*TortVmax*) venular tortuosity. Central Retinal Artery and Vein Equivalent (*CRAE, CRVE*) and the Arteriole-to-Venule ratio (*AVR*) qualify vessel calibers and were measured in a zone 2 to 3 optic disc radii from the center of the optic disc. Among the eight parameters, *TortA, TortAmax, TortV* and *TortVmax* mean values were normalized by natural log transformation for association analysis.

##### Replication Cohorts

Standard retinal fundus images using digital fundus camera from 1091 individuals from LBC1936 were collected at the recruitment stage and three years later, retinal traits were measured at a subsequent wave of testing using SIVA v3.1[32,33] (Singapore I Vessels Assessment), at a mean age of 72.5 years (SD 0.7). A total of 897 and 976 individual’s retinal fundus images centered between the macula and optic disc from Croatia-Korčula and Croatia-Split cohorts were collected using digital fundus camera and retinal traits were quantified using SIVA v3.1[32,33]. SIVA is a semi-automated software which can be used to measure the retinal vascular parameters including retinal vascular tortuosity and vascular caliber from retinal images. After automatic detection of the optic disc, it placed a grid with reference to the center of the optic disc. Then the tortuous vessels were identified and tortuosity traits including *TortA*, and *TortV* were measured using the standard grading protocol by the software; this process was monitored by trained graders and adjusted manually if necessary.

### Genotyping, quality control and imputation

#### Discovery cohorts

GoDARTS samples were genotyped using the Affymetrix 6.0 (n=927) and Illumina Human Omni Express (n=809) platforms. The poor quality variants, samples were excluded based on the quality control (QC) criteria included the following: SNPs call rate < 95%, Hardy–Weinberg equilibrium (HWE) P value < 10^−6^, sample call rate < 95%, sample relatedness (IBD >0.8), and mismatch between reported and genotypic gender information. QC’d genotype data were imputed using IMPUTE2[47,48] on the basis of 1000 Genome Projects reference panel for all population. Finally, ancestry information of the individuals was derived using EIGENSTRAT[49] and first three principal components (PCs) were used for the association analyses to adjust the population stratification. ORCADES samples were genotyped with either the Illumina HumanHap300 bead chip (n=890) or the Illumina Omni1 (n=304) or Illumina Omni Express bead chips (n=1073). Alleles were called in Bead Studio/Genome Studio (Hap300/Omni) using Illumina cluster files. Subjects were excluded if they fulfilled any of the following criteria: genotypic call rate <98%, mismatch between reported and genotypic sex, unexpectedly low genomic sharing with first or second degree relatives, excess autosomal heterozygosity, or outliers identified by IBS clustering analysis. We excluded SNPs on the basis of minor allele frequency (<0.01/monomorphism), HWE (P<10^−6^), call rate (<97%). Given the very high overlap in SNPs between the two Omni chips, the intersection of QC’d SNPs was used to impute and phase individuals’ genotyped on the Omni arrays together, whilst the Hap300 individuals were phased and imputed, separately. Samples were phased using Shapeit v2[50]. Imputation was carried out using IMPUTE2 and the 1,000 genomes reference panel. All ancestries phase1 integrated v3 reference panel, with a secondary reference panel of local exome sequences, sequenced using the Agilent Sure Select All Exon Kit v2.0 and Illumina 100 bp paired end reads (average 30x depth), derived from 90 ORCADES subjects chosen to optimally represent the haplotypes present. Imputations for the Hap300 and Omni subjects were then combined to form a combined panel of 37.5m SNPs for 2222 subjects[51]. Imputed genotypes for 658, 1078, 1358 individuals from the GoDARTS dataset 1, GoDARTS dataset 2 and ORCADES cohorts, respectively, were used for the three independent GWAS analysis.

#### Replication Cohorts

LBC1936 samples were genotyped at the Wellcome Trust Clinical Research Facility, Edinburgh, using the Illumina Human 610Quad BeadChip. Individuals were excluded based on unresolved gender discrepancy, relatedness, call rate (≤0.95), and evidence of non-Caucasian descent. SNPs were included if they met the following conditions: call rate ≥ 0.98, minor allele frequency ≥ 0.01, and Hardy-Weinberg equilibrium test with P ≥ 0.001. Imputation to the 1000 Genomes (March 2012 release) reference set was performed using minimac software. A total of 1398 participants from the two independent Croatian replication cohorts were available for the analysis and subjects were genotyped on different genotyping platforms including Illumina CNV370v1 and CNV370-Quadv3 for Croatia-Korčula (n=378), and Illumina CNV370-Quadv3 and IlluminaOmniExpressExome-8v1_A for Croatia-Split (n=376). Samples and markers were excluded based on the following QC metrics; SNPs call rate < 98%, HWE with P value < 10^−6^, sample call rate < 97%, MAF < 1%, outliers identified by IBS clustering analysis and unresolved gender discrepancy. Imputation was carried out using IMPUTE2 software and 1000G Phase I v3 (March 14, 2012) reference panel.

#### Statistical analyses

We performed association analyses with each data sets from GoDARTS separately for each of the eight retinal traits using SNPTEST V2.5[47], linear regression assuming an additive genetic model, adjusting for 3 ancestry PCs, age at eye examination and gender. Subsequently, markers with low imputation quality scores (< 0.4) and minor allele frequency cutoffs (< 0.03) were filtered from each GWAS summary output data separately. Then we performed the meta-analysis using a fixed-effects model in GWAMA[52] with the QC filtered data sets. Association analysis in ORCADES was performed for each of the eight retinal traits, using linear mixed modelling to account for relatedness and assuming an additive genetic model, adjusting for 3 ancestry PCs, age at eye examination and gender, using MMscore in ProbABEL[53]. As in GoDARTS, markers with low imputation quality scores (< 0.4) and minor allele frequency cutoffs (< 0.03) were filtered and meta-analysis was performed with the GoDARTS and ORCADES results using GWAMA. The strand alignment and build check between studies were performed prior to meta-analysis. Also, the genomic inflation factor (λ) was estimated by GWAMA (λ=0.99). All statistical analyses and QCs were performed using SNPTEST v2.5[47], ProbABEL[53], GWAMA[52], PLINK v1.09[54], EIGENSTRAT[49], custom shell scripts, and R scripts. Manhattan plots, Quantile-Quantile plots and forest plots were generated using in-build R scripts, and metafor - R package[55]. Regional plots were generated using the Locus Zoom tool[56] and other data processing was performed using R scripts. Conditional analyses were performed in SNPTEST v2.5 using the genome-wide associated loci in the *COL4A2* region, conditioned on lead SNPs (rs56399312). Also, this new locus was conditioned on previously reported genome-wide significant SNPs (rs4773144, rs11617955, rs9515203) associated with coronary artery disease (CAD).

#### Sensitivity analyses

We performed an association test with an additive model adjusted for age, gender, and first three principal components in the diabetes cohort (GoDARTS) using 759 samples without any cardiovascular events prior to the retinal screening date.

#### Replication-analyses

The top three SNPs (P ≤ 1.07×10^−07^) near *ACTN4, TMEM132D*, and *COL4A2* from the discovery stage for the tortuosity traits were taken forward for examination in three replication cohorts of European ancestry. In the LBC1936 cohort, association analysis was performed for arterial and venular tortuosity traits using linear regression model adjusting for age at eye examination, sex, and 3 ancestry PCs, using mach2qtl. Similarly, in the Croatia - Split, - Korčula cohorts, association analysis were performed for each traits separately using the mixed model in R - hglm package to account for kinship derived using gkin function of the GenABEL package[57].

Then we combined the summary association statistics for lead SNPs associated with *TortA* and *TortV* from the two discovery and three replication cohorts and effect estimates from each cohort were presented in the forest plots using metafor - R package. Due to the difference in the units of the beta and standard errors between the discovery and replication studies arising from different approaches to measurement we standardized the effect estimates (using Cohen’s d) from each of the individual study cohort.

#### Power calculation

The statistical power of detecting SNPs associations with the quantitative traits in two stage GWAS was calculated using the GWASPower/QT.

#### *In-silico* look-ups of the novel variants for clinical outcomes

We performed *in-silico* look-ups of variants of interest for cardiovascular related outcomes including coronary artery disease, myocardial infarction, hypertension, HDL, and triglycerides from the CARDIoGRAMplus C4D consortium[20], GLGC [21] and the ICBPGWAS analysis[22]. The CARDIoGRAMplusC4D 1000 Genomes-based meta-analysis data comprised of 60,801 CAD cases and 123,504 controls from European, South Asian, and East Asian descent. In the GLGC, genetic data from 188,577 individuals of European, East Asian, South Asian, and African ancestry were used to examine the genetic loci associated with blood lipids levels. The ICBP GWAS investigated the genetic loci associated with systolic and diastolic blood pressure traits in 200,000 individuals of European descent. We retrieved summary association results for the index SNPs from these studies to investigate the association of the lead SNPs for *TortA*, and *TortV* with cardiovascular outcomes.

A recent study reported the association of *ACTN4* locus with heart rate[44]. In order to examine whether the lead SNPs associated with *TortV* in *ACTN4* were also associated with heart rate, we checked the LD (r^2^>0.8) between our SNPs and the index SNP (rs11083475) for heart rate in that study. Furthermore, we investigated the association of these SNPs with pulse rate in the UK Biobank data. This data comprised of 112,008 participants who had a measure of pulse rate at the main interview and had genotype data. We extracted the imputed genotypes for these SNPs from the interim release data set of the UK Biobank[23] and performed multiple linear regressions including covariates of age, gender, and the first ten principal components obtained using EIGENSTRAT.

#### *In-silico* functional annotation

The sentinel genome-wide significant variants were mapped to the gene, 20 kb upstream/downstream using BEDTools[58], and UCSC Genome Browser[59]. Top SNPs were queried in the HaploReg v4.1 database[60] to catalogue the all SNPs near noncoding variants with r^2^ > 0.8, and RegulomeDB[61], and GWAS catalog databases[62] used to explore the known and predicted regulatory elements and relevant genetic association studies. Functional effects of the top genes were predicted using the Encyclopedia of DNA Elements[43] (ENCODE) project and Roadmap Epigenomics projects which aggregate the information about the transcription factor, motifs, histone modification, and chromatin states. Additionally, functional elements were investigated using HaploReg, UCSC Genome Browser, and RegulomeDB. We used the expression Quantitative Trait Loci (eQTL) browser database in Genotype-Tissue Expression[41] (GTEx) to examine the cis-eQTLs for the top retinal traits associated SNPs mapped to the gene within the genomic region. Gene Visible web database from GENEINVESTIGATOR which integrates manually curated gene expression data from microarray and RNAseq experiments, was used to find the expression level of the genes, associated with tortuosity traits, in the human tissues.

## Acknowledgements

We are grateful to all the participants in the GoDARTS study, the general practitioners, the Scottish School of Primary Care for their help in recruiting the participants, and to the whole team, which includes interviewers, computer and laboratory technicians, clerical workers, research scientists, volunteers, managers, receptionists, and nurses. The study complies with the Declaration of Helsinki. We acknowledge the support of the Health Informatics Centre, University of Dundee for managing and supplying the anonymized data and NHS Tayside, the original data owner. The Wellcome Trust United Kingdom Type 2 Diabetes Case Control Collection (GoDARTS) was funded by The Wellcome Trust (072960/Z/03/Z, 084726/Z/08/Z, 084727/Z/08/Z, 085475/Z/08/Z, 085475/B/08/Z) and as part of the EU IMI-SUMMIT program.

ORCADES was supported by the Chief Scientist Office of the Scottish Government (CZB/4/276, CZB/4/710), the Royal Society, the MRC Human Genetics Unit, Arthritis Research UK and the European Union framework program 6 EUROSPAN project (contract no. LSHG-CT-2006-018947). DNA extractions were performed at the Wellcome Trust Clinical Research Facility in Edinburgh. We would like to acknowledge the invaluable contributions of the research nurses in Orkney, the administrative team in Edinburgh and the people of Orkney.

We thank the Lothian Birth Cohort 1936 (LBC1936) participants and team members who contributed to these studies. Phenotype collection was supported by Age UK (The Disconnected Mind project). Genotyping was funded by the BBSRC (BB/F019394/1). The work was undertaken by The University of Edinburgh Centre for Cognitive Ageing and Cognitive Epidemiology, part of the cross council Lifelong Health and Wellbeing Initiative (MR/K026992/1). Funding from the BBSRC and Medical Research Council (MRC) is gratefully acknowledged.

The Croatia-Korčula and Croatia-Split study were funded by grants from the Medical Research Council (UK), European Commission Framework 6 project EUROSPAN (Contract No. LSHG-CT-2006-018947), European Commission Framework 7 project BBMRI-LPC (FP7 313010), the Republic of Croatia Ministry of Science, Education and Sports research grant (216-1080315-0302) and the Croatian Science Foundation (grant 8875). We would like to acknowledge the staff of several institutions in Croatia that supported the field work, including but not limited to the University of Split and Zagreb Medical Schools and Croatian Institute for Public Health. The SNP genotyping for the Korčula cohort was performed in Helmholtz Zentrum München, Neuherberg, Germany.

VAMPIRE team: Parts of the VAMPIRE software and its use for measuring the image set described here was funded by the Leverhulme Trust project RPG-419 “Discovery of retinal biomarkers for genetics with large cross-linked data sets”. VAMPIRE 3.1 has been developed under funding from EPSRC (EPSRC EP/M005976/1), the EU (REVAMMAD ITN).

For the analysis of the association of the identified genetic variants with heart rate, this research has been conducted using the UK Biobank Resource under Application Number 20405.

## Supporting Information

**S1 Fig. Retinal fundus image.** Solid lines (red for arterioles and dark blue for venules) represent the vessels detected automatically and measured by VAMPIRE (Vasculature Assessment and Measurement Platform for Images of the REtina) software (version 3.0, Universities of Edinburgh and Dundee, UK). Dotted lines (light blue) represent the measurement zones on a fundus image; based on optic disc (light blue circle) location and radius.

**S2 Fig.** Meta-analysis on genome-wide association results from two independent discovery cohorts. Manhattan plots for six quantitative retinal traits. A. represents the results from Optic Disc Radius (*ODradius*), B. represents the results from retinal arteriolar tortuosity maximum (*TortAmax*), C. represents the results from retinal venular tortuosity (*TortVmax*), D. represents the results from Central Retinal Arteriolar Equivalent (*CRAE*), E. represents the results from Central Retinal Venular Equivalent (*CRVE*), and F. represents the results from Arteriole-to-Venule ratio (*AVR*). The blue and red horizontal lines indicate the suggestive and genome-wide significance threshold (P<5×10^−8^), respectively.

**S3 Fig.** Quantile-quantile plots of GoDARTS-ORCADES meta-analysis for eight quantitative retinal blood vessel traits. Shaded areas represent 95% confidence intervals. Naturally log transformed -TortA: retinal arteriolar tortuosity, TortAmax: maximum retinal arteriolar tortuosity, TortV: retinal venular tortuosity, TortVmax: maximum retinal arteriolar tortuosity. ODradius: Optic Disc Radius, CRAE: Central Retinal Arteriolar Equivalent, CRVE: Central Retinal Venular Equivalent, AVR: Arteriole-to-Venule ratio.

**S4 Fig.** Regional association plots of index SNP reached p-value < 5×10^−7^ in the metaanalysis of the two discovery study cohorts (GoDARTS and ORCADES). a) *TortA* b) maximum *TortA* c-d) *TortV* e) *ODradius* f) maximum *TortV*.

**S5 Fig.** Box plots represent the expression level of genes associated with retinal blood vessel traits. a) *COL4A2* b) *COL4A1* c) *ACTN4* d) *CAPN12*. These plots were created using GENEVESTIGATOR which integrates manually curated gene expression data from microarray and RNAseq experiments. Blue lines at the bottom of the box indicates the gene expression level across 451 tissues in human. Y-axis depicts the top ten tissues and the sample size shown in secondary Y-axis. Colour scale (Low to High) at the top depicts the gene expression range in log2scale.

**S6 Fig.** eQTL annotation of top GWAS variants for quantitative retinal vessel traits. Y-axis represents tissue-specific gene expression data which is normalized by rank normalization method while x-axis shows the GWAS lead variant genotypes. a) TortA associated SNP, rs7991229 is correlated with *COL4A2* expression in heart left ventricle; b) TortA associated SNP, rs9515212 is correlated with *COL4A2* expression in heart left ventricle tissue; c) TortV associated SNP, rs1808382 is correlated with *CAPN12* expression in artery aorta; d) TortV associated SNP, rs1808382 is correlated with *ACTN4* expression in artery aorta.

**S7 Fig.** Conditional analysis of the genome-wide significant variant (rs56399312) at *COL4A2* locus. Locus zoom plots for the *COL4A2* locus associated region (GoDARTS) conditioned on the CAD associated SNPs (rs11617955, rs4773144, rs9515203) reported previously in the GWAS study. a) Top SNP (TortA) Conditioned on rs11617955, b) Top SNP (TortA) Conditioned on rs4773144 c) Top SNP (TortA) Conditioned on rs9515203.

**S8 Fig.** Multi-tissue eQTL comparison for TortV associated SNP, rs1808382 is correlated with a) *ACTN4* expression and b) *CAPN12* expression.

**S9 Fig.** Locus zoom plot for the *ACTN4* locus associated with TortV also associated with pulse rate in UK Biobank. The lead SNP, rs1808382 associated with TortV (Discovery and replication stage) in that region is indicated by purple colour solid diamond.

**S1 Table**. Significant SNPs for each quantitative retinal vascular traits that reached P<7×10^−7^ in the meta-analysis of discovery cohorts.

**S2 Table**. SNPs in *COL4A2* conditioned on top SNP rs56399312 (discovery cohorts), associated with *TortA*. TortA, retinal arteriolar tortuosity.

**S3 Table**. Summary of previously reported significant SNPs associated with Optic Disc area, CRAE, and CRVE look-ups in GoDARTS-ORCADES meta-analysis study.

**S4 Table**. *Insilico* functional annotation of significant SNPs for quantitative retinal vascular traits.

**S5 Table**. Significant top hits that reached P<1×10^−7^ for quantitative retinal vascular traits as eQTLs (GTEx) in different tissues.

**S6 Table**. Top SNPs in *COL4A2* associated with TortA conditioned on reported coronary artery disease SNPs.

**S7 Table**. Summary of significant SNPs (P<8×10^−07^) associated with tortuosity traits from discovery stage, replicated in myocardial infarction (MI) and coronary artery disease (CAD) GWAS.

**S8 Table**. Summary of previously reported significant SNPs associated with coronary artery disease (CARDioGramplus C4D) look-ups in GoDARTS-ORCADES meta-analysis study for retinal arteriolar tortuosity trait.

**S9 Table**. Genome-wide significant SNPs (P<8×10^−07^) for retinal tortuosity traits from discovery stage, are associated with different traits (Type2Diabetes Knowledge Portal and ICBP).

**S10 Table**. Sensitivity analyses using GoDARTS (diabetes).

**S11 Table**. Lead SNPs associated with TortV are also associated with heart rate in UK Biobank.

## Author Contributions

The study was designed by C.NA.P, A.SF.D, and E.T for GoDARTS cohort, J.F.W for ORCADES cohort, I.J.D for LBC1936 cohort, O.P for Croatia-Split, and Croatia-Korcula cohort. VAMPIRE software was designed and developed by E.T, T.M, D.R, E.B, S.K.V, and B.D. Retinal images were collected and analysis was performed by E.T, T.M, J.F.W, L.B, M.K, D.R, V.V, and H.C. Genotype data processing and statistical analysis was conducted by A.V, K.E.S, P.K.J, L.B, M.K, S.H, V.V, C.H and K.Z. Bioinformatics analysis was performed by A.V. The manuscript was drafted by A.V, C.NA.P A.SF.D, and revised by E.T, J.F.W, T.M, I.J.D, S.H, E.R.P and K.Z. All the authors reviewed the manuscript.

